# Integrated bioinformatics and single-cell analysis identifies vascular aging-related hub genes and immune drivers in atherosclerosis

**DOI:** 10.64898/2026.04.14.718580

**Authors:** Xiaofeng Chen, Kunlin Zhou, Weichun Wang, Jingying Wu

**Affiliations:** Department of Cardiology, Second Affiliated Hospital of Fujian Medical University, Quanzhou, Fujian Province, 362000, China

**Keywords:** atherosclerosis, vascular aging, immune infiltration, cell senescence, RNA-sequencing, single-cell RNA sequencing

## Abstract

Atherosclerosis (AS) is a chronic inflammatory disease closely linked to vascular senescence, yet the specific molecular mechanisms connecting aging processes to AS pathogenesis remain incompletely understood. This study integrated transcriptomic data from GEO datasets (GSE100927 and GSE43292) to identify vascular aging-related differentially expressed genes (VARDEGs). Following batch effect correction, 28 VARDEGs were screened and subjected to functional enrichment, protein-protein interaction (PPI) network analysis, and immune infiltration assessment. Seven hub genes (*MMP9, APOE, TNF, ICAM1, PPARG, CYBA,* and *NCF2*) were identified and experimentally validated via qRT-PCR, confirming their significant upregulation in AS samples. Receiver operating characteristic (ROC) analysis demonstrated high diagnostic accuracy for six of these genes (AUC > 0.7), with *TNF* exhibiting superior performance. Immune infiltration analysis revealed profound alterations in 28 immune cell types, particularly monocytes and T cells, which correlated strongly with hub gene expression. Furthermore, single-cell RNA sequencing analysis (GSE184073) localized the expression of core genes predominantly to monocytes and T cells, highlighting *TNF* overexpression in T cells as a potential critical driver. Finally, molecular docking simulations suggested that curcumin exhibits strong binding affinity to these hub genes, particularly *PPARG*, providing a mechanistic basis for its therapeutic potential. Collectively, this study elucidates the landscape of vascular aging-related genes in AS, identifies novel diagnostic biomarkers, and proposes potential therapeutic targets involving immune modulation and natural compounds.

## Introduction

Atherosclerosis, a chronic cardiovascular disease, is the leading cause of death in humans and is the underlying cause of peripheral vascular disease, coronary heart disease, and stroke [1]. Its pathogenesis is complex and involves multiple cell types, including endothelial cells (ECs), vascular smooth muscle cells (VSMCs), adventitia fibroblasts, macrophages, and other immune cells [2, 3]. The key factors for the development of atherosclerosis include endothelial dysfunction, leukocyte adhesion, macrophage foam formation, and changes in the VSMC phenotype [4, 5]. Atherosclerosis is driven by both traditional and non-traditional risk factors [6]. Hypertension, diabetes, smoking, and hypercholesterolemia, which lead to the dysfunction of Ecs and VSMCs, have been identified as the traditional risk factors for atherosclerosis [7, 8]. Meanwhile, the nontraditional risk factors, including C-reactive protein, oxidative stress, and inflammation, accelerate arteriosclerosis [9]. Despite aggressive control of the risk factors for atherosclerosis, many people still die from its acute complications. The incidence and mortality rates of atherosclerosis increase significantly with age. Several studies have demonstrated the important role of vascular aging in the development of atherosclerosis [10, 11].

Although various epidemiological and clinical studies have demonstrated an intimate connection between vascular aging and atherosclerosis, the underlying mechanisms should be thoroughly examined. Vascular aging is a specific type of aging that involves the senescence of various cells in the arteries. Atherosclerosis is a chronic inflammatory disease in which cellular senescence occurs in atherosclerotic plaques [12]. Senescent cells secrete inflammatory cytokines and chemokines through the senescence-associated secretory phenotype (SASP), which induces local and systemic inflammatory responses, immune system activation, tissue damage, fibrosis, apoptosis, and dysfunction [13]. The SASP of senescent cells has several proatherogenic effects that may involve vascular remodeling, plaque formation, and rupture [2, 12]. These events lead to vascular aging and accelerate the development of atherosclerosis via a series of structural and functional changes. Therefore, focusing on the role of vascular aging in the pathogenesis of atherosclerosis is crucial for delaying atherosclerosis. However, the potential role of vascular aging in the diagnosis, prognosis, and treatment of atherosclerosis remains unclear.

In this study, we integrated an atherosclerosis dataset from the Gene Expression Omnibus (GEO) and vascular aging-related genes (VARGs) obtained from GeneCards to identify reliable vascular aging-related differentially expressed genes (VARDEGs) in atherosclerosis. Through rigorous bioinformatics analysis, we identified 28 VARDEGs and constructed a protein–protein interaction (PPI) network to screen for key hub genes. We further validated the expression of seven hub genes (*MMP9, APOE, TNF, ICAM1, PPARG, CYBA,* and *NCF2*) via qRT-PCR and evaluated their diagnostic potential using receiver operating characteristic (ROC) curve analysis. Moreover, we employed immune infiltration analysis and single-cell RNA sequencing (scRNA-seq) to dissect the relationship between these hub genes and the immune microenvironment, revealing a strong association with monocyte and T cell dynamics. Finally, molecular docking simulations were performed to explore the therapeutic potential of curcumin in targeting these aging-related hubs. Our findings provide novel insights into the molecular crosstalk between vascular aging and atherosclerosis, identifying potential biomarkers for early diagnosis and new targets for therapeutic intervention.

## Materials and methods

### Animals

Male apoE^−/−^mice (6-weeks-old) were purchased from Gempharmatech Co., Ltd (Jiangsu, China). The protocols of animal experiments were performed following the National Institutes of Health Laboratory Animal Care and Use Guidelines and approved by the Animal Ethics Committee at Fujian Medical University. ApoE^−/−^ mice were randomized into the following two groups: Control, AS. ApoE^−/−^ mice were fed a high-fat diet containing 0.15% cholesterol and 21% fat for 12 weeks to establish the animal model of atherosclerosis. ApoE^−/−^mice fed a standard chow diet served as the control.

### Total RNA extraction and quantitative reverse transcription polymerase chain reaction (qRT-PCR)

Total RNA was extracted from mouse tissues using TRIzol reagent (Sangon Biotech, China). cDNA was synthesized using the High-Capacity cDNA Reverse Transcription Kit (TIANGEN Biotech, China). qRT-PCR was performed on a StepOne Plus system (Applied Biosystems) using SuperReal PreMix Plus (SYBR Green) (TIANGEN). Gene expression levels were normalized to GAPDH using the (2–ΔΔCt) method. The primer sequences were as follows: GAPDH (B661304, Sangon Biotech, Shanghai, China); MMP9, 5′-CAGACCAAGGGTACAGCCTG-3′ (forward) and 5′-ATACAGCGGGTACATGAGCG-3′ (reverse); APOE, 5′-AAGATGAAGGCTCTGTGGGC-3′ (forward) and 5′-AATCCCAGAAGCGGTTCAGG-3′ (reverse); TNF, 5′-ATGTCTCAGCCTCTTCTCATTC-3′ (forward) and 5′-GCTTGTCACTCGAATTTTGAGA-3′ (reverse); ICAM1, 5′-TAATGTCTCCGAGGCCAGGA-3′ (forward) and 5′-CGAGCTTCAGAGGCAGGAAA-3′ (reverse); PPARG, 5′-AAGCCGTGCAAGAGATCACA-3′ (forward) and 5′-TGGTCATGAATCCTTGGCCC-3′ (reverse); CYBA, 5′-ACCATCAAGCAGCCACCTAC-3′ (forward) and 5′-TCAAGCAGGAGCCACTGAAG-3′ (reverse); and NCF, 5′-CAGCCCTTTCTGTCCCTCAG-3′ (forward) and 5′-CGCATGCCTTTAATCCCAGC-3′ (reverse).

### Downloaded data

We used the R package GEOquery [14] from the GEO database [15] (https://www.ncbi.nlm.nih.gov/geo/) to download the atherosclerosis datasets GSE100927 [16] and GSE43292 [17]. The samples of datasets GSE100927 and GSE43292 were all from *Homo sapiens*, and the tissue source was the artery (Table 1). The chip platform of the dataset GSE100927 was GPL17077, which included 69 atherosclerosis and 35 control samples. The chip platform of the dataset GSE43292 was GPL6244, which contained 32 atherosclerosis and 32 control samples. All atherosclerosis and control samples were included in this study. The GeneCards database [18] (https://www.genecards.org/) is a collection of VARGs and provides comprehensive information on human genes. We used the term “Vascular Aging” as a search keyword and only kept “Protein Coding” VARGs, and a total of 69 VARGs were obtained (S1 Table).

**Table 1.**
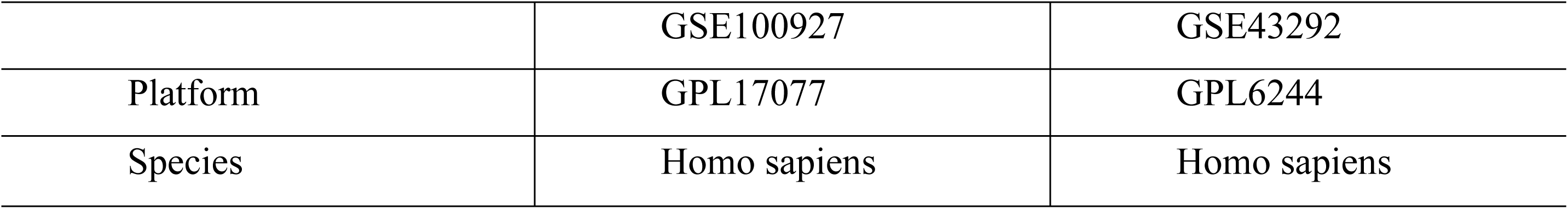
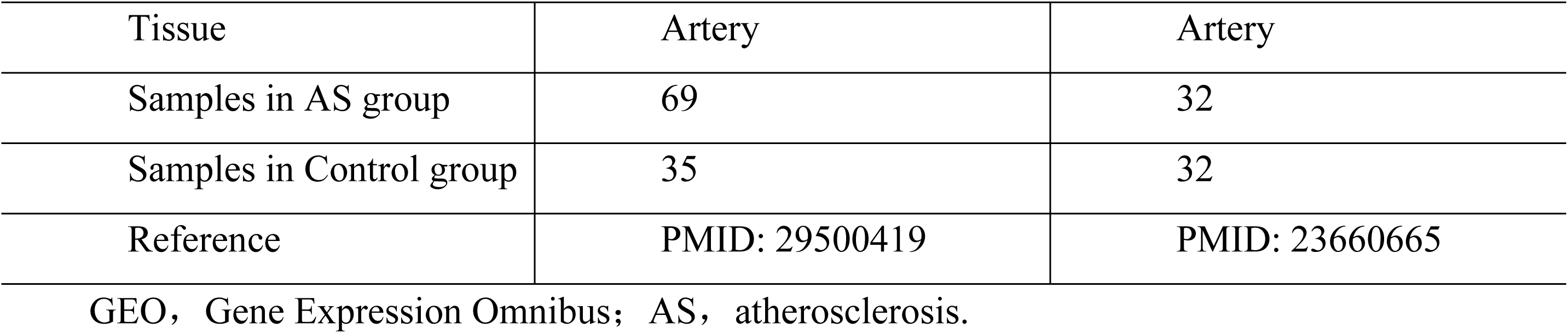
GEO microarray chip information.

|  | GSE100927 | GSE43292 |
| --- | --- | --- |
| Platform | GPL17077 | GPL6244 |
| Species | Homo sapiens | Homo sapiens |
| Tissue | Artery | Artery |
| Samples in AS group | 69 | 32 |
| Samples in Control group | 35 | 32 |
| Reference | PMID: 29500419 | PMID: 23660665 |

GEO, Gene Expression Omnibus; AS, atherosclerosis.

The R package sva [19] was used to debatch GSE100927 and GSE43292 to obtain the combined GEO datasets. The combined dataset included 101 atherosclerotic and 67 control datasets. Finally, the R package limma [20] was used to standardize the combined GEO datasets, annotate the probes, and normalize them. The expression matrices before and after removing the batch effect were subjected to a principal component analysis (PCA) [21] to verify the effect of removing the batch effect.

### DEGs related to atherosclerosis related vascular aging

Based on the sample grouping of the combined GEO datasets, the samples were divided into the atherosclerosis and control groups. Differential analysis of genes between the atherosclerosis and control groups was performed using the R package limma, setting | logFC | > 0.25 and adj. *P* < 0.05 for the DEG threshold. DEGs with logFC > 0.25 and adj. *P* < 0.05 were considered upregulated, whereas those with logFC < −0.25 and adj. *P* < 0.05 were downregulated. The *P*-value correction method was Benjamini–Hochberg (BH). The results of the difference analysis were plotted as volcano plots using the R package ggplot2. To obtain VARDEGs, all DEGs with |logFC| > 0.25 and adj. *P* < 0.05 in the integrated GEO dataset were drawn Venn diagram with VARGs. Furthermore, heat maps were drawn using the R package pheatmap and the R package Rcircos map chromosomal location [22].

### Gene ontology (GO) and Kyoto Encyclopedia of Genes and Genomes (KEGG) pathway enrichment analysis

GO analysis [23] is commonly used for large-scale functional enrichment studies, including biological processes (BP), cell components (CC), and molecular functions (MF). KEGG [24] is a widely used database storing information on genomes, biological pathways, diseases, and drugs. We used the R package clusterProfiler [25] to perform GO and KEGG pathway enrichment analyses of VARDEGs. The item screening criteria were adj. *P* < 0.05 and false discovery rate (FDR) (q value) < 0.25 were considered significant. The *P*-value correction method was BH.

### Gene set enrichment analysis (GSEA)

GSEA [26] was used to evaluate the distribution trend of genes in a predefined gene set in a gene table ranked by correlation with the phenotype and thus determine their contribution to the phenotype. In this study, genes from the combined GEO datasets were first ranked by logFC values, and the R package clusterProfiler was used to perform GSEA on all genes in the integrated GEO datasets (combined datasets). Regarding the parameters used in the GSEA, the number of seeds and computations was 2020 and 1000, respectively. The minimum and maximum number of genes contained in each gene set was 10 and 500, respectively. The Molecular Signatures Database [27] (https://www.gsea-msigdb.org/gsea/msigdb) was used to access c2 gene sets. Cp. All. V2022.1. Hs. Symbols. GMT [All Canonical Pathways] (3050) was used for GSEA. Significance in the screening criteria for GSEA was set at adj. *P* < 0.05 and FDR (q value) < 0.25. The *P*-value correction method was BH.

### PPI network and hub gene screening

The PPI network is composed of proteins interacting with each other and participating in biological signaling, gene expression regulation, all aspects of life processes, such as energy and substance metabolism, and cell cycle regulation. A systematic analysis of the interaction of proteins in biological systems is crucial for understanding the working principle of proteins in biological systems, the reaction mechanism of biological signals, energy and substance metabolism under special physiological conditions, such as diseases, and the functional relationship between proteins. The STRING database [28] (https://string-db.org/) was searched to determine interactions between known and predicted proteins. In this study, the STRING database was used based on VARDEGs, with a minimum interaction coefficient of >0.700 (minimum required interaction score: high confidence, 0.700) used as the standard to construct the PPI network related to VARDEGs. The closely connected local regions in the PPI network may represent molecular complexes with specific biological functions. Genes that interacted with other genes in the PPI network were selected for subsequent analyses. Five algorithms in the CytoHubba software [29] were applied as follows: maximal clique centrality (MCC), degree, maximum neighborhood component (MNC), edge percolated component (EPC), and closeness [30]. In the PPI network, the VARDEG scores were calculated, and then the TOP10 VARDEGs were selected according to the scores. Finally, the genes obtained by the five different algorithms were interfaced, and a Venn diagram was drawn for analysis. The intersecting genes in the algorithms were vascular age-related hub genes.

### Construction of regulatory network

Transcription factors (TFs) control gene expression by interacting with hub genes at the post-transcriptional stage. The ChIPBase database [31] (http://rna.sysu.edu.cn/chipbase/) was searched to retrieve TFs, and only TFs with the sum of “number of samples found (upstream)” and “Number of samples found (downstream)” > 6 were retained. The regulatory effects of TFs on hub genes were analyzed, and the mRNA–TF regulatory network was visualized using the Cytoscape software.

Additionally, miRNAs play important regulatory roles in biological development and evolution. They regulate various target genes, and the same target gene can be regulated by multiple miRNAs. To analyze the hub genes and the relationship between miRNA, through the StarBase v2.0 database [32], only miRNAs with “pancancerNum > 6” were retained, and the mRNA–miRNA regulatory network was visualized using the Cytoscape software.

### Differential expression verification and ROC curve analysis

To further explore the expression differences of hub genes between atherosclerotic and control samples in the combined GEO datasets, a group comparison map was drawn based on the expression levels of hub genes. Next, the R package pROC was used to plot the ROC curve of the hub genes and calculate the area under the curve (AUC). We evaluated the diagnostic effect of hub gene expression on atherosclerosis occurrence. The AUC of the ROC curves were generally between 0.5 and 1. The closer the AUC is to 1, the better the diagnostic performance. AUC values of 0.5–0.7, 0.7–0.9, and >0.9 were considered low, moderate, and high, respectively.

### Immune infiltration analysis

Single-sample gene set enrichment analysis (ssGSEA) [33] quantifies the relative abundance of each immune cell infiltrate. First, each infiltrating immune cell type was labeled, including activated CD8+ T cells, activated dendritic cells, gamma delta T cells, and natural killer cells. Human immune cells have various subtypes, such as regulatory T cells. Subsequently, enrichment scores calculated using the ssGSEA were used to represent the relative abundance of each immune cell infiltrate in each sample, resulting in an immune cell infiltration matrix for the combined GEO datasets. Subsequently, the R package ggplot2 was used to draw group comparison maps to show the expression differences in immune cells between the control and atherosclerosis groups of the combined GEO datasets. Immune cells with significant differences between the two groups were screened for subsequent analyses. The correlation between immune cells was calculated using the Spearman algorithm, and the R package pheatmap was used to draw a correlation heatmap to show the correlation analysis results of the immune cells themselves. The correlation between hub genes and immune cells was calculated based on the Spearman algorithm, and the correlation bubble plot was drawn using the R package ggplot2.

### Single-cell RNA sequencing (scRNA-seq) analysis

The raw single-cell RNA-seq data from the GSE184073 dataset were downloaded and processed for subsequent analysis. Quality control and data preprocessing were performed using the Seurat R package (v4.1.0). Low-quality cells were filtered out based on the following criteria: cells with fewer than 200 or more than 7500 detected genes, as well as cells with a mitochondrial content exceeding 20%. Read counts were normalized per cell using a scale factor of 10,000 and log-transformed. The data were then scaled using the ScaleData function in Seurat to regress out potential confounding sources of variation. The top 2,000 highly variable genes were selected for dimensionality reduction via principal component analysis (PCA). The top 30 principal components were used for cell clustering and visualization. Cell clusters were identified using the FindNeighbors and FindClusters functions (resolution = 0.05) in Seurat. Cell type annotation was performed by combining marker gene expression analysis via the FindAllMarkers function and automated cell type labeling using the SingleR package. Finally, the expression of hub geens across cell clusters and between sample groups was visualized using the FeaturePlot and DotPlot functions in Seurat.

### Molecular docking

The binding affinity and interaction mode between the candidate drug and the target were analyzed using AutodockVina 1.2.2. The molecular structures were obtained from the PubChem compound database (https://pubchem.ncbi.nlm.nih.gov/). The three-dimensional coordinates of hub genes (*MMP9, APOE, TNF, ICAM1, PPARG, CYBA,* and *NCF2*) were downloaded from the PDB (http://www.rcsb.org/pdb/home/home.do). For the docking analysis, all protein and molecular files were converted to the PDBQT format, all water molecules were excluded, and polar hydrogen atoms were added. The grid box was centered on covering the structure domain of each protein and accommodating free molecular movement. The molecular docking study was conducted through Autodock Vina 1.2.2 (http://autodock.scripps.edu/).

### Statistical analysis

All data processing and analyses in this study were based on the R software (version 4.3.0). Regarding the comparison of continuous variables between the two groups, normally distributed variables were compared using the independent Student’s t-test, whereas differences in non-normally distributed variables were analyzed using the Mann–Whitney U test (Wilcoxon rank-sum test). The Kruskal–Wallis test was used to compare three or more groups. Spearman’s correlation analysis was performed to calculate the correlation coefficients between the different molecules. All two-sided *P* < 0.05 were considered significant.

## Results

### Technology roadmap

S1 Fig. presents a flowchart of the comprehensive analysis of VARDEGs. We used the online GEO microarray expression profiling datasets GSE100927 and GSE43292 for research. Differentially expressed VARGs were analyzed using correlation analysis, GO enrichment analysis, KEGG pathway enrichment analysis, GSEA, and PPI network analysis. Bioinformatics analysis was performed to identify hub genes. Transcription factor-hub gene regulatory networks, miRNA-hub gene regulatory networks, and immune cell infiltration analyses were performed. We then determined the diagnostic significance of hub genes using ROC curve analysis.

### Merging of atherosclerosis datasets

First, the R package sva was used to perform batch-effect removal on the atherosclerosis datasets GSE100927 and GSE43292 to obtain the combined GEO datasets. Subsequently, distribution boxplots (S2 Fig. A, B) were used to compare the expression values of the datasets before and after removing the batch effect. Second, a PCA plot (S2 Fig. C, D) was used to compare the distribution of low-dimensional features before and after batch effect removal. The results of the distribution boxplot and PCA plot show that the batch effect of the samples in the atherosclerosis dataset was eliminated after batch removal.

### DEGs related to atherosclerosis-related vascular senescence

Data from the combined GEO datasets were divided into the atherosclerosis and control groups. To analyze the differences in gene expression values between the two groups in the combined GEO datasets, the R package limma was used for differential analysis of the combined GEO datasets to obtain DEGs between the two datasets. The results were as follows: integrated GEO dataset (combined datasets) a total of 3559 met the | logFC | > 0.25 and adj. *P* < 0.05 threshold of DEGs. Under the threshold, up expressed genes (logFC > 0.25 and adj., *P* < 0.05), a total of 1818 down expressed genes (logFC < −0.25 and adj. *P* < 0.05), a total of 1741, according to the variance analysis results of the data set map volcano (Fig. 1A).

**Fig. 1.**
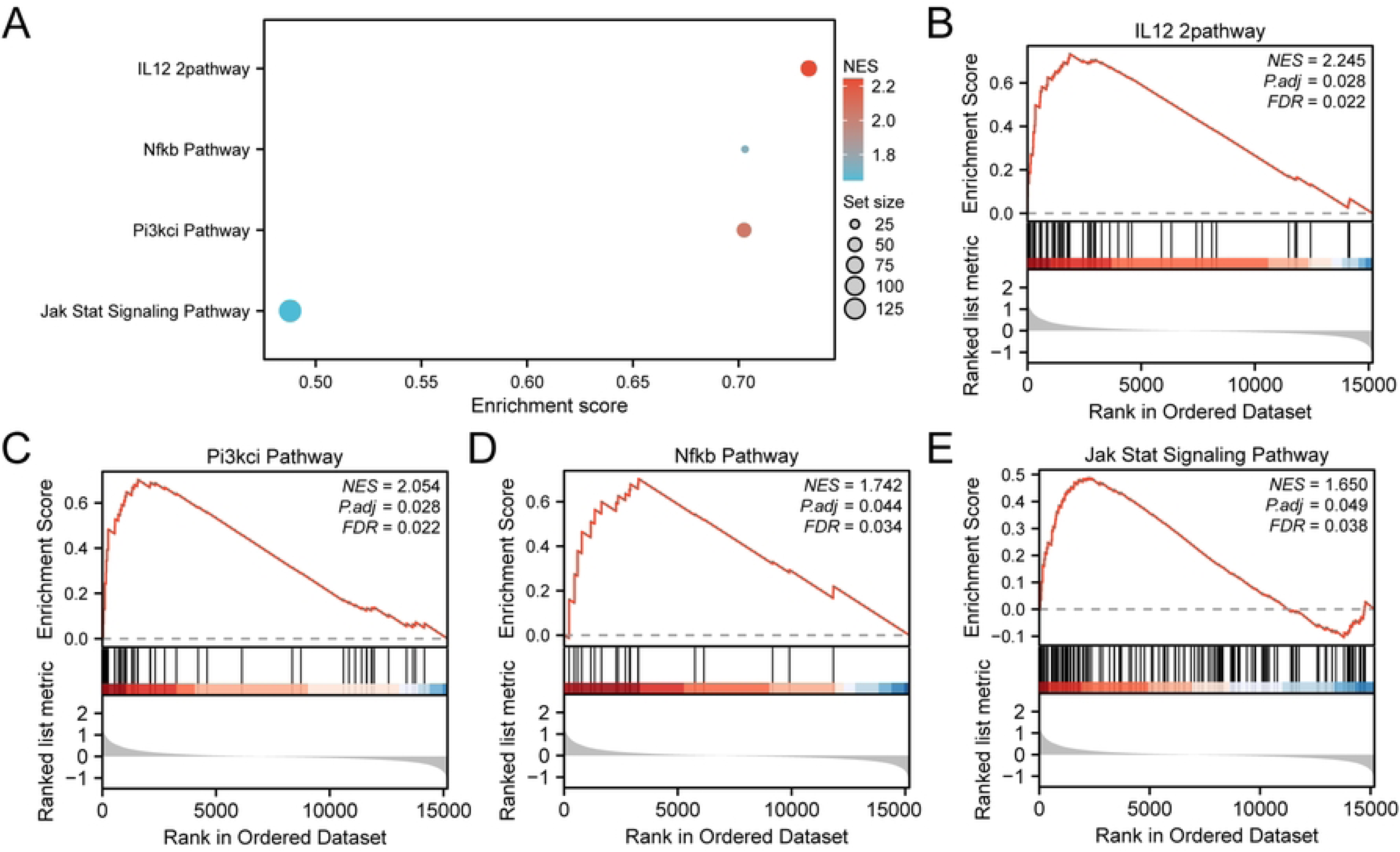
Differential gene expression analysis. A. Volcano plot of differentially expressed genes in atherosclerosis and control samples in the combined GEO datasets. B. DEGs and VARGs Venn diagrams in the combined GEO datasets. C. Heat map of VARDEGs associated with vascular aging in in the combined datasets. D. Chromosomal mapping of VARDEGs. In the heat map grouping, green represents the control sample, and purple indicates the atherosclerosis sample. In the heat map, red represents a high expression, and blue indicates a low expression. DEGs, differentially expressed genes; VARDEGs, vascular age-related differentially expressed genes.

For VARDEGs meeting the threshold | logFC | > 0.25 and adj. *P* < 0.05, a Venn diagram was plotted to demonstrate the intersection between DEGs and VARGs (Fig. 1B). A total of 28 VARDEGs were identified (S2 Table). According to the intersection results, the logFC-ranked TOP10 upregulated and TOP10 downregulated VARDEGs between different sample groups in the datasets were analyzed, and the R package pheatmap was used to draw a heatmap to display the analysis results (Fig. 1C). Finally, the locations of the 28 VARDEGs on the human chromosomes were analyzed using the R package Rcircos to construct a chromosome localization map (Fig. 1D). Chromosome mapping showed that more VARDEGs were located in chromosomes 1 and 2 as follows: *ZMPSTE24*, *NCF2*, *CHI3L1*, and *NLRP3* were located in chromosome 1, and *APOB, DPP4*, *PDE1A*, and *MYO1B* were located in chromosome 2.

### GO and KEGG enrichment analyses

Gene Ontology (GO) and Kyoto Encyclopedia of Genes and Genomes (KEGG) pathway analyses were used to further explore the relationships between BP, CC, MF, and biological pathway (KEGG) of 28 VARDEGs and atherosclerosis. The 28 VARDEGs were used for GO and KEGG pathway enrichment analyses. The results showed that 28 VARDEGs were mainly enriched in muscle cell proliferation (including smooth muscles) and regulation, as well as EC proliferation (BP) and its regulation (S5 Table); NADPH oxidase complex, endocytic vesicle, membrane microdomain, membrane raft, neuronal cell body(CC); Superoxide-generating NAD(P)H oxidase activity, oxidoreductase activity, acting on NAD(P)H, oxygen as acceptor, 3′,5′-cyclic-GMP phosphodiesterase activity, 3′,5′-cyclic-AMP phosphodiesterase activity and oxidoreductase activity, acting on NAD(P)H and other molecular functions (MF). It was also enriched in lipid and atherosclerosis, fluid shear stress, atherosclerosis, the AGE–RAGE signaling pathway in diabetic complications, leukocyte transendothelial migration, and the osteoclast differentiation pathway (KEGG). The results of the GO and KEGG pathway enrichment analyses are shown in the bar diagram (Fig. 2A) and bubble diagram (Fig. 2B).

**Fig. 2.**
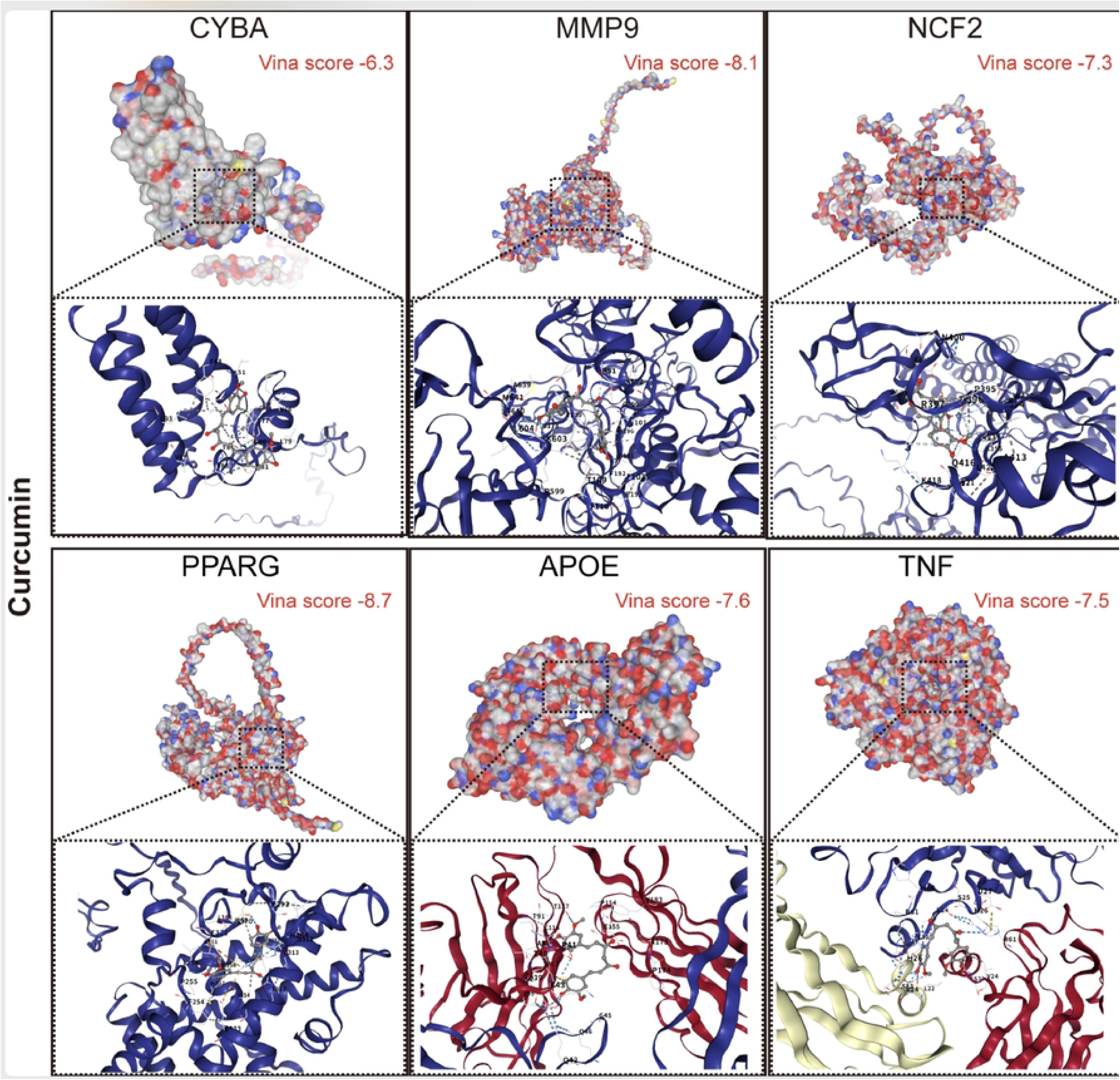
GO and KEGG enrichment analysis for VARDEGs. A, B. Results of the GO and KEGG pathway enrichment analyses of VARDEGs. Bar graph (A) and bubble plot (B) show the BP, CC, MF, and biological pathway (KEGG). GO terms and KEGG terms are shown on the ordinate. C–F. Results of the GO and pathway (KEGG) enrichment analyses of the VARDEG network diagram showing BP (C), CC (D), MF €, and KEGG (F). Green nodes represent items, purple nodes indicate molecules, and the lines represent the relationship between the items and molecules. VARDEGs, vascular aging-related differentially expressed genes; GO, gene ontology; KEGG, Kyoto Encyclopedia of Genes and Genomes; BP, biological process; CC, cellular component; MF, molecular function. The bubble size in the bubble plot represents the number of genes, and the color of the bubble represents the size of the adj. P-value, the redder the color, the smaller the adj. P-value, and the bluer the color, the larger the adj. P-value. The screening criteria for the GO and pathway (KEGG) enrichment analyses were significance at adj. *P* < 0.05 and FDR (q value) < 0.25. The Benjamini–Hochberg method was used for correcting P-values.

Meanwhile, the network diagram of BP, CC, MF, and biological pathway (KEGG) was drawn according to the GO and pathway (KEGG) enrichment analyses (Fig. 2C–F). The lines show the corresponding molecules and annotations of the corresponding entries. The larger the nodes, the more molecules the entries contain. More genes were enriched in the regulation of SMC proliferation, SMC proliferation, muscle cell proliferation (BP), and lipids and atherosclerosis (KEGG).

### GSEA for atherosclerosis

GSEA was performed to investigate the expression of all genes in the combined GEO datasets, the biological processes involved, and the link between the cellular components affected and molecular functions (Fig. 3A). All genes in the combined GEO datasets were significantly enriched in the IL12 2pathway (Fig. 3B), Pi3kci pathway (Fig. 3C), Nfkb pathway (Fig. 3D), JAK–STAT signaling pathway (Fig. 3E), and other biologically relevant functions and signaling pathways (S6 Table).

**Fig. 3.**
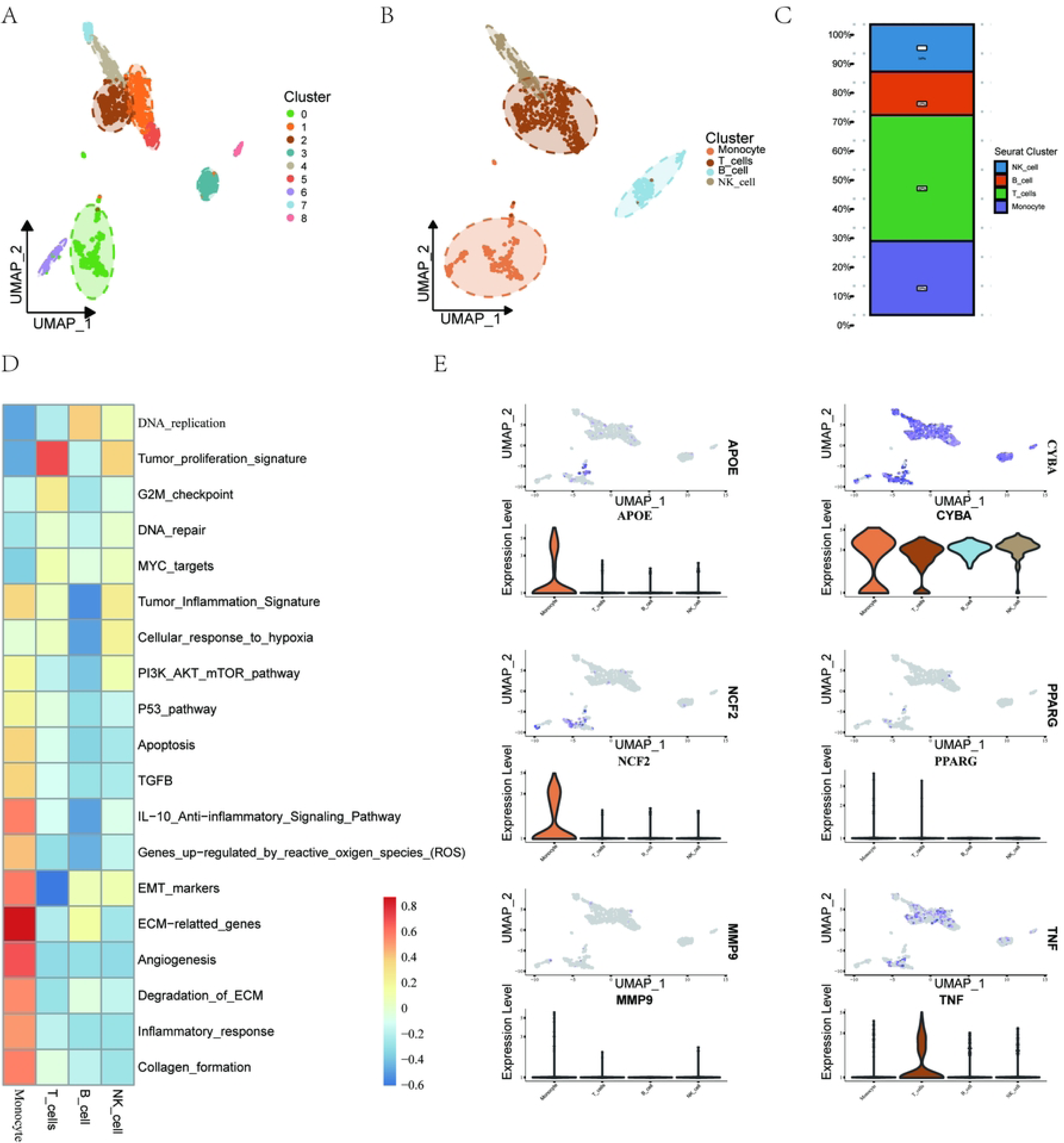
GSEA for combined datasets. A. GSEA 4 biological function bubble plot of integrated GEO datasets. B–E. GSEA showed that all genes were significantly enriched in the IL12 2pathway (B), Pi3kci Pathway (C), Nfkb pathway (D), and JAK–STAT signaling pathway €. In the bubble plot, the size and color of the bubble represents the number of enriched genes and size of the NES value, respectively. The redder the color, the greater the NES value, and the bluer the color, the smaller the NES value. The screening criteria for the GSEA were significance at adj. *P* < 0.05 and FDR (q value) < 0.25. The Benjamini–Hochberg method was used for correcting *P*-values. GSEA, gene set enrichment analysis, NES, normalized enrichment score.

### Construction of a PPI network and screening of hub genes

A PPI network of 28 VARDEGs was constructed using the STRING database (Fig. 4A). The results of the PPI network showed that 18 VARDEGs were related: *APOB, APOE, FTO, PPARG, MMP9, ICAM1, TNF, BMP2, CDKN2A, DNMT1, CYBA, NOX4, NCF2, MFGE8, KDR, NLRP3, PDE1A,* and *PDE1C*.

**Fig. 4.**
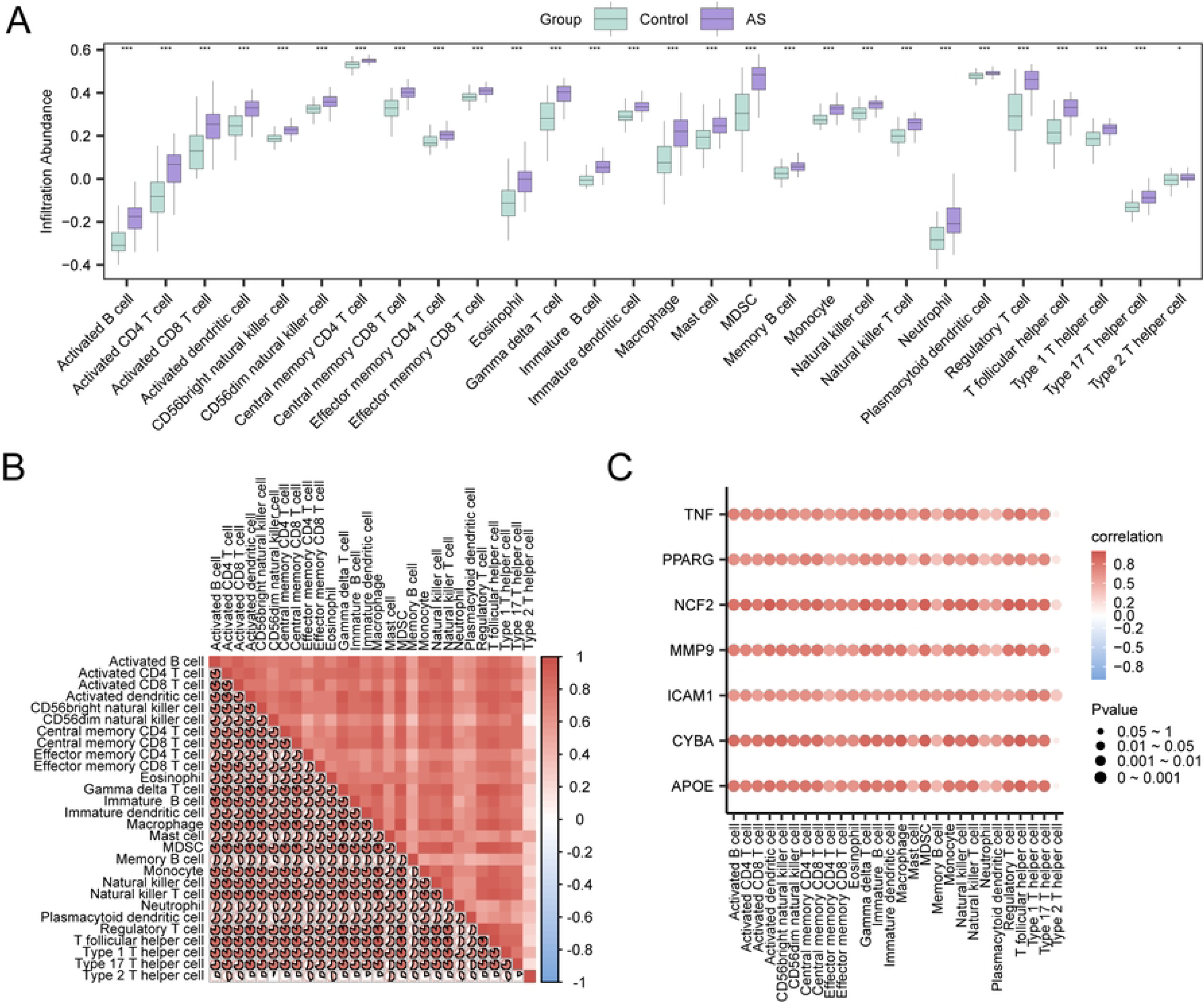
PPI network and hub gene analysis. A. PPI network of VARDEGs calculated from the STRING database. B–F. Top 10 VARDEGs PPI network calculated by five algorithms of the CytoHubba plug-in, including MCC (B), MNC (C), Degree (D), EPC (E), and closeness (F). G. Venn diagram of the top 10 VARDEGs calculated by five algorithms of the CytoHubba plugin. PPI, protein–protein interaction; VARDEGs, vascular aging-related differentially expressed genes; MCC, maximal clique centrality; MNC, maximum neighborhood component; EPC, edge percolated component.

Subsequently, the five algorithms of the CytoHubba plug-in of the Cytoscape software were used to calculate the scores of 18 VARDEGs, which were ranked according to the scores. The five algorithms were MCC, degree, MNC, EPC, and closeness. Then, the TOP10 VARDEGs from the five algorithms were used to draw PPI networks: MCC (Fig. 4B), degree (Fig. 4C), MNC (Fig. 4D), EPC (Fig. 4E), and closeness (Fig. 4F). The color of the circles from red to yellow represents the scores from high to low. Finally, the intersection of the genes of the five algorithms was measured, and a Venn diagram (Fig. 4G) was drawn for analysis. The intersecting genes from these algorithms were identified as hub genes: *MMP9*, *APOE*, *TNF*, *ICAM1*, *PPARG*, *CYBA*, and *NCF2*.

### Construction of regulatory networks

TFs that bind to hub genes were obtained from the ChIPBase database, and the mRNA-TF regulatory network was constructed and visualized using the Cytoscape software (Fig. 5A). Among them, six were hub genes and 42 were TFs (S3 Table).

**Fig. 5.**
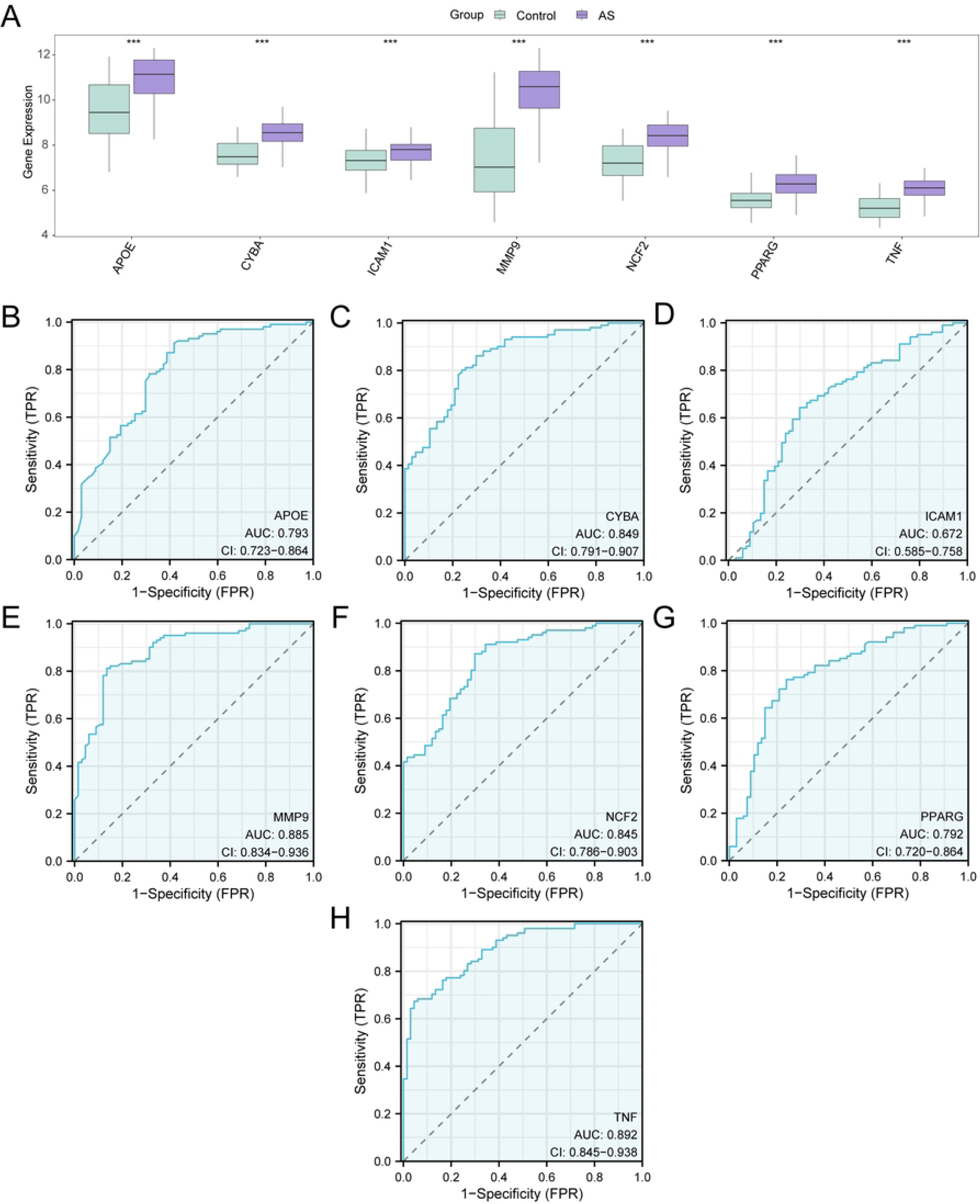
Regulatory network of hub genes. A. mRNA–TF regulatory network of hub genes. B. mRNA–miRNA regulatory network of hub genes. MRNAs are shown in yellow, transcription factors in blue, and miRNAs in red. TF: transcription factor. C-I. Validation of hub genes by qRT-PCR. \*\**P* <0.01, \*\*\**P* <0.001,\*\*\*\**P* < 0.0001 (n =6 for each group).

Then, miRNAs related to hub genes were obtained from the TarBase database, and the mRNA-miRNA regulatory network was constructed and visualized using the Cytoscape software (Fig. 5B). Among them, there were 3 Hub Genes and 30 miRNAs, and the specific information is shown in S4 Table.

### Validation of the 7 hub genes using qRT-PCR

To confirm the 7 hub genes identified using the aforementioned analyses and to determine the variations of expression at the mRNA level, qRT-PCR were performed (Fig. 5C–I). The results revealed that the 7 hub genes were significantly upregulated in AS group when compared with the control group (*P*<0.01), which was concordant with the results obtained from the bioinformatics analysis.

### Differential expression verification and ROC curve analysis

To explore the expression differences of hub genes in the Combined GEO Datasets, the group comparison figure (Fig. 6A) shows the differential analysis results of the expression levels of seven hub genes (*MMP9*, *APOE*, *TNF*, *ICAM1*, *PPARG*, *CYBA*, and *NCF2*) in atherosclerosis and control samples in the combined GEO datasets. The differential results showed that the expression levels of seven hub genes in atherosclerosis and control samples of the combined GEO datasets were highly significant (*P* < 0.001). Next, we used the R package pROC to draw ROC curves based on the expression of hub genes in the combined GEO datasets (Fig. 6B–H). The ROC curve showed that the expression levels of the six hub genes (*MMP9*, *APOE*, *TNF*, *PPARG*, *CYBA*, and *NCF2*) in atherosclerotic samples showed a certain accuracy with different groups (0.7 < AUC < 0.9). The expression level of the hub gene *ICAM1* in atherosclerosis samples showed a low accuracy among different groups (0.5 < AUC < 0.7).

**Fig. 6.**
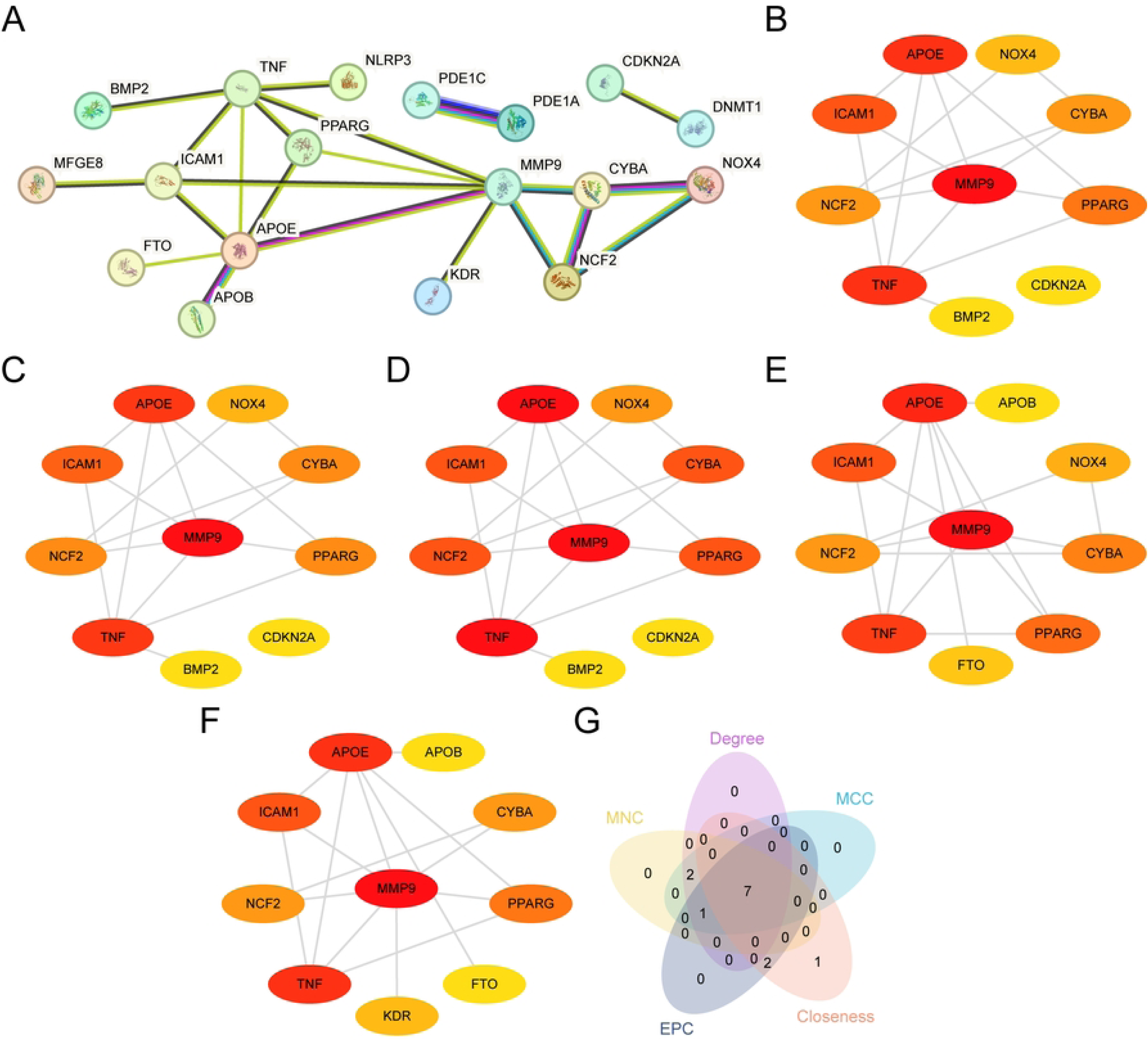
Differential expression validation and ROC curve analysis. A. Group comparison plots of hub genes in atherosclerosis and control samples from combined GEO datasets. B–I. ROC curves of hub genes (hub genes) APOE (B), CYBA (C), ICAM1 (D), MMP9 (E), NCF2 (F), PPARG (G), and TNF (H) in integrated GEO datasets. Group comparison plots show the control samples in yellow and atherosclerosis samples in purple. When AUC was 0.7–0.9, it had some accuracy, and when AUC was 0.5–0.7, it had low accuracy. *** indicates high significance at P < 0.001. AS, atherosclerosis; ROC, receiver operating characteristic; AUC, area under the curve; TPR, true positive rate; FPR, false positive rate.

### Immune infiltration analysis of atherosclerosis

The expression matrices of the combined datasets were used to calculate the immune infiltration abundance of 28 immune cells using the ssGSEA algorithm. First, differences in the expression of infiltrating immune cells in different groups are shown by group comparison plots. The group comparison diagram (Fig. 7A) showed that all the 28 immune cells were statistically significant (*P* < 0.05), namely activated B cell, activated CD4+ T cell, activated CD8+ T cell, activated dendritic cell, CD56bright natural killer cell, CD56dim natural killer cell, central memory CD4+ T cell, central memory CD8+ T cell, effector memory CD4+ T cell, effector memory CD8+ T cell, eosinophil, gamma-delta T cell, immature B cell, immature dendritic cell, macrophage, mast cell, myeloid-derived suppressor cell (MDSC), memory B cell, monocyte, natural killer cell, natural killer T cell, neutrophil, plasmacytoid dendritic cell, regulatory T cell, T follicular helper cell, type 1 T helper cell, type 17 T helper cell, and type 2 T helper cell. A correlation heatmap was used to show the correlation results of 28 immune cell infiltration abundances in the immune infiltration analysis of the combined GEO datasets (Fig. 7B). Most immune cells had strong positive correlations, and immune cell macrophages and MDSC had the most significant positive correlation (r = 0.967, *P* < 0.05). Finally, the correlation between hub genes and abundance of immune cell infiltration is demonstrated using a correlation bubble plot (Fig. 7C). Most immune cells showed a strong positive correlation, and the gene NCF2 and immune cell-activated dendritic cells had the strongest significant positive correlation (r = 0.945, *P* < 0.05).

**Fig. 7.**
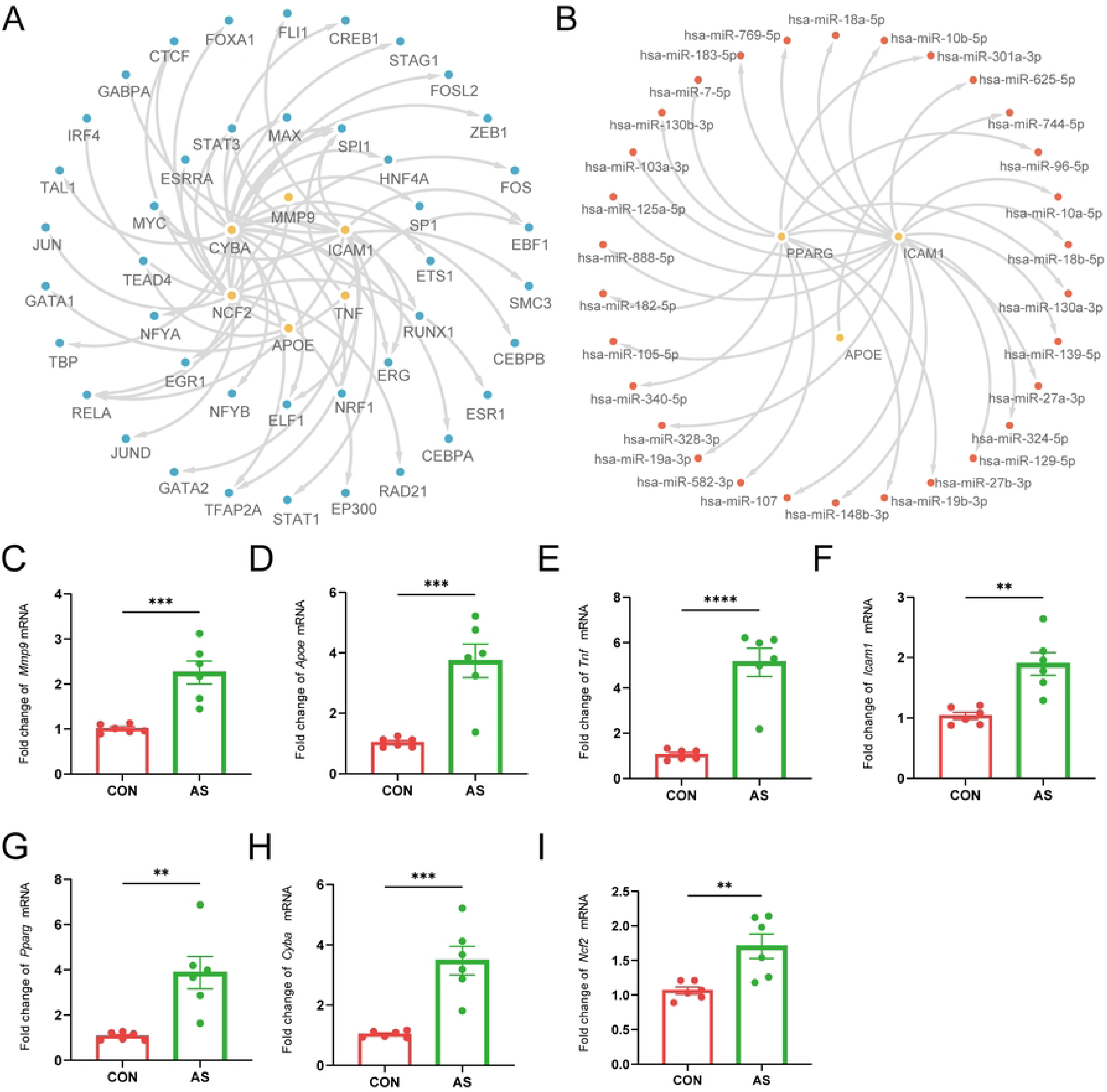
Immune infiltration analysis by ssGSEA algorithm. A. Group comparison plots of immune cells in atherosclerosis and control samples from the combined GEO datasets. B. Heat map of immune cell infiltration abundance in the integrated GEO datasets (combined datasets). C. Bubble plot of the correlation between hub genes and immune cell infiltration abundance in the integrated GEO datasets (combined datasets). ssGSEA, single-sample gene-set enrichment analysis. Group comparison plots show control samples in green and atherosclerosis samples in purple. The absolute values of correlation coefficients (r value) <0.3, 0.3–0.5, 0.5-0.8, and >0.8 indicate no, weak, moderate, and correlations, respectively. In the correlation heat map, red indicates a positive correlation, and blue represents a negative correlation. The depth of the color represents the strength of the correlation. AS, atherosclerosis. * represents *P* < 0.05, indicating statistical significance; *** represents *P* < 0.001, indicating a high statistical significance.

### Single-cell validation of hub genes expression and immune infiltration analysis

Next, we analyzed the single-cell RNA sequencing dataset of coronary artery disease (GSE184073). Cell subpopulations were annotated into four major types—NK cells, T cells, B cells, and monocytes—based on canonical marker genes and automated annotation using SingleR(Fig. 8A-B). Among these, T cells constituted the largest proportion, followed by monocytes, suggesting that these two cell types may play pivotal roles in the pathogenesis of atherosclerosis(Fig. 8C). Projection of the six core genes onto the single-cell data revealed their predominant expression in monocytes and T cells(Fig. 8E). Notably, our previous analyses identified TNF as exhibiting the highest ROC performance, with significant upregulation observed specifically in T cells. This finding implies that elevated TNF expression within monocytes (and potentially T cells) may serve as a critical driver of disease progression. These results are consistent with our immune infiltration analysis.

**Fig. 8.**
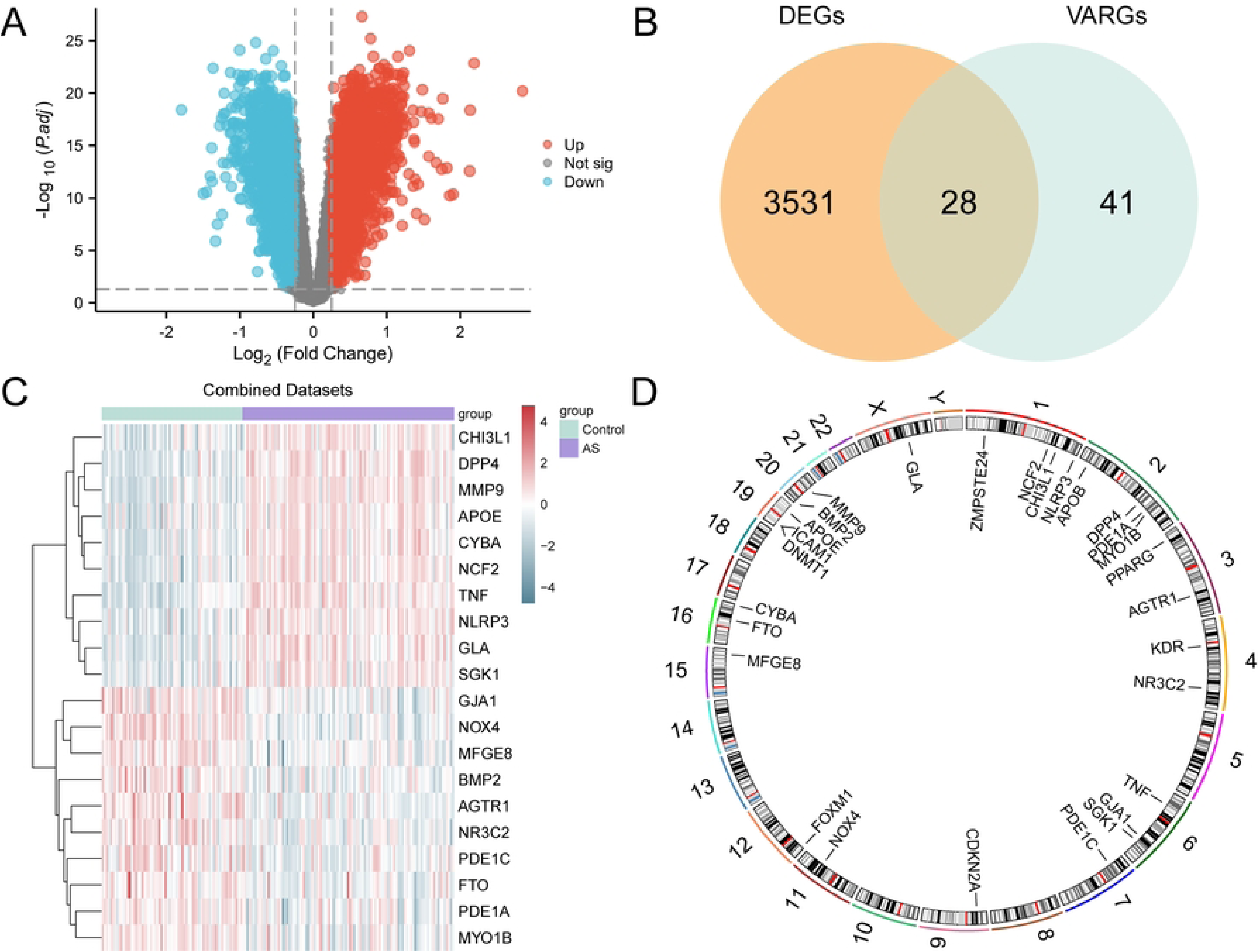
Single-cell transcriptomic landscape of the atherosclerotic immune microenvironment (GSE184073). A. UMAP visualization of eight unsupervised cell clusters. B. Cell type annotation into four major lineages (T cells, Monocytes, B cells, NK cells) based on marker genes and SingleR. C. Proportional composition of cell types, highlighting the dominance of T cells and Monocytes. D. Distinct metabolic pathway activity scores across the four annotated cell populations. E. Projection of the six hub genes, revealing their predominant expression in Monocytes and T cells, with *TNF* specifically upregulated in T cells.

### Validation of curcumin-VARDEGs interaction by molecular docking

Curcumin has been demonstrated to possess anti-atherosclerotic properties[34], primarily attributed to its anti-inflammatory effects; however, the precise mechanisms underlying these effects remain inadequately understood. In this study, the authors explored the interaction between curcumin and six key genes—*MMP9*, *APOE*, *TNF*, *PPARG*, *CYBA*, and *NCF2*—utilizing molecular docking analysis(Fig. 9). The findings revealed that curcumin exhibited significant binding affinity with all six target genes, with binding energies consistently below -5 kcal/mol. Notably, the binding affinity with PPARG was the most pronounced. This molecular docking analysis substantiates the interaction between curcumin and these hub genes at a molecular level, suggesting that curcumin may exert its anti-atherosclerotic effects by mitigating aging processes.

**Fig. 9.**
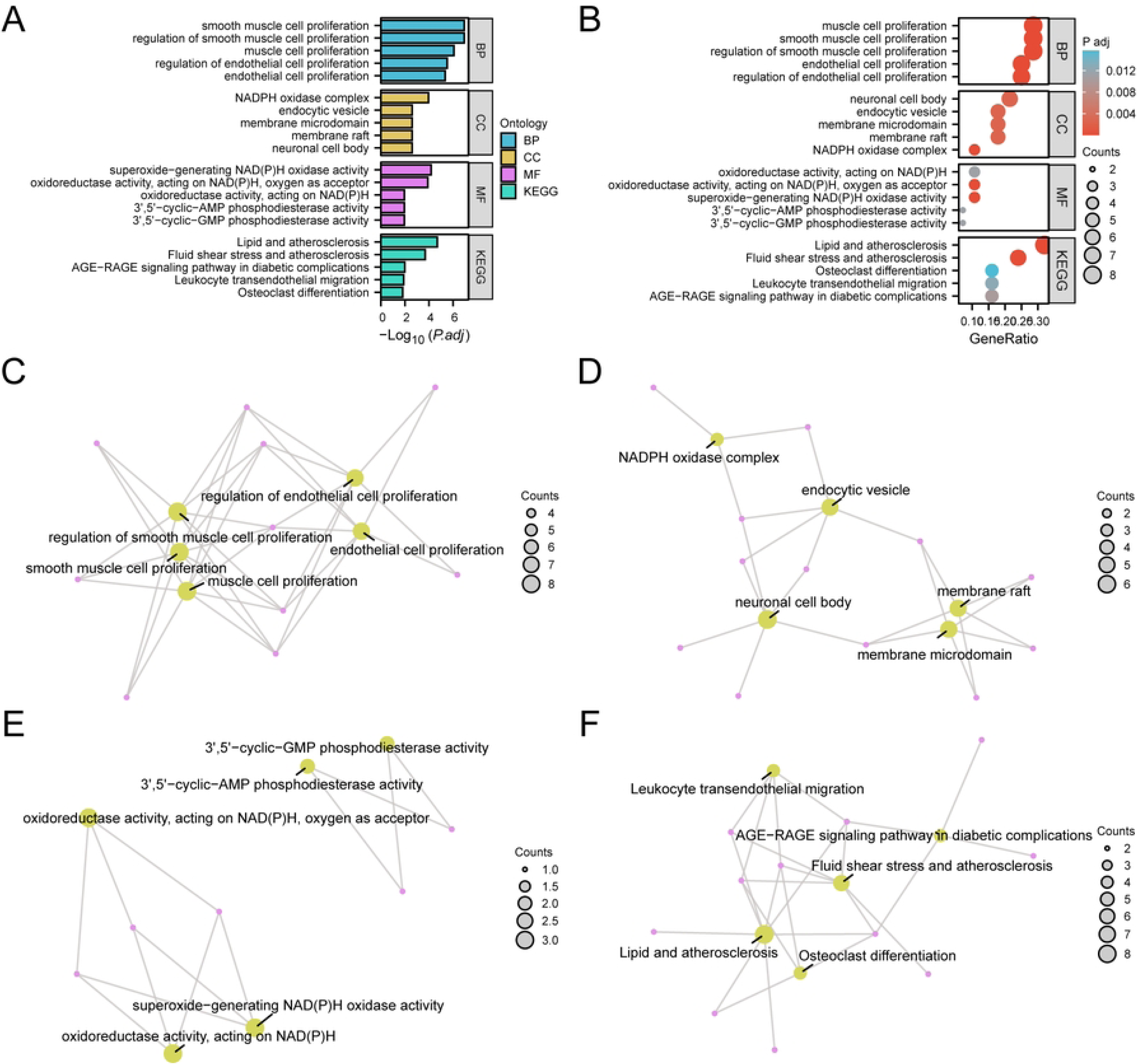
Molecular docking analysis illustrating the binding interactions and affinities between curcumin and the six hub genes (*MMP9, APOE, TNF, PPARG, CYBA,* and *NCF2*).

## Discussion

Atherosclerosis is the leading cause of cardiovascular death worldwide [35]. Atherosclerotic plaque rupture leading to thrombosis may lead to life-threatening clinical complications including myocardial infarction, stroke, and sudden death [36]. However, the mechanisms underlying atherosclerosis remain unclear. Vascular aging is a key factor in the development and progression of atherosclerosis. Vascular remodeling related to vascular aging can lead to patients with high sensitivity to cardiovascular risk factors (such as hyperlipidemia, hypertension, diabetes, and smoking) who are more susceptible to atherosclerosis [37].

Upon exposure to cardiovascular risk factors, senescent cells accumulate in the arteries and release excessive inflammatory factors through the SASP, which may activate the senescence pathway and further senesce neighboring cells [38]. Inflammation is an important biological marker, and its activation can lead to high platelet reactivity, leading to blood platelet coagulation potential, increased blood adhesion, and thrombosis [39]. These events lead to vascular aging and accelerate the development of arteriosclerosis via a series of structural and functional changes. Vascular aging and arteriosclerosis interact in a vicious circle, which jointly promotes the occurrence and development of vascular-related diseases, such as coronary heart disease, hypertension, cerebrovascular disease, and chronic kidney disease[38, 40]. The impairment of endothelial function or barrier integrity caused by vascular aging can accelerate the deposition of LDL in the intimal layer, potentially triggering platelet aggregation and thrombosis [2]. The vascular inflammatory response affects the recruitment and adhesion of inflammatory cells and promotes the formation of atherosclerotic plaques [39, 41]. Thus, many forms of vascular aging are involved in the development and progression of arteriosclerosis. However, the specific mechanisms remain unclear, and further studies are needed to broaden our knowledge of vascular aging in the pathogenesis of arteriosclerosis.

To the best of our knowledge, we identified 28 potential vascular age-related genes in arteriosclerosis for the first time using bioinformatics analysis. We further identified seven hub genes associated with arteriosclerosis, including *MMP9*, *APOE*, *TNF*, *ICAM1*, *PPARG*, *CYBA*, and *NCF2*, using PPI network and key module analyses. The roles of these genes in arteriosclerosis have been studied extensively. For example, *MMP9* is involved in plaque formation by degrading the extracellular matrix to weaken arterial walls, leading to cardiovascular remodeling [42]. *APOE* participates in the transformation and metabolism of lipoproteins, and *APOE* knockout mice are the most common models of atherosclerosis. *TNF* and *ICAM1* are genes that regulate inflammatory cytokines and participate in the chronic inflammatory process of arteries, playing an important role in vascular aging and arteriosclerosis, which further indicates that inflammation may be a same mechanism of vascular aging and arteriosclerosis[43, 44]. *PPARG* is a gene related to lipid metabolism[45]. *PPARG* increases insulin sensitivity, promotes lipogenesis, and plays an anti-inflammatory and anti-atherosclerotic role[46]. Both *CYBA* and *NCF2* genes are involved in the metabolism of NAPDH oxidase. NADPH oxidase is the main source of reactive oxygen species (ROS), and oxidative stress is a risk factor for atherosclerosis[47, 48]. However, the roles of these vascular senescence-related genes in the regulation of atherosclerosis have not been fully explored [1].

GO analysis showed that 28 VARDEGs were enriched in muscle cell proliferation (including smooth muscles) and its regulation as well as EC muscle cell proliferation and its regulation (BP). They were also enriched in lipids, atherosclerosis, fluid shear stress, the AGE–RAGE signaling pathway in diabetic complications, leukocyte transendothelial migration, and the osteoclast differentiation pathway (KEGG). Atherosclerosis involves the pathological activation of various cell types, including immune cells (e.g., macrophages and T cells), SMCs, and ECs [49]. Increasing evidence has suggested that the transition of SMCs to other cell types, known as phenotypic switching, plays an important role in the development and complications of atherosclerosis [50]. All genes in the combined GEO datasets were significantly enriched in the IL12 [51], Pi3kci [52], Nfkb [53] and JAK–STAT signaling pathways [54], which are related to cell senescence and vascular aging. Inflammation is a key driver of atherosclerosis, and the upstream source of inflammatory factors might be cellular senescence [39]. Cell senescence promotes the occurrence and development of atherosclerosis by secreting proinflammatory cytokines (including IL-12, etc.) through the SASP [38]. The JAK–STAT pathway is an important signaling pathway that regulates the initiation and progression of atherosclerosis, and JAK–STAT activation has been observed in atherosclerotic lesions [55]. Further studies have reported that inhibition of the JAK–STAT signaling pathway can attenuate cell senescence and the SASP [56]. Therefore, treatments targeting the JAK–STAT signaling pathway may alleviate atherosclerosis by delaying cell senescence.

Next, we constructed the mRNA–TF and mRNA–miRNA regulatory networks of hub genes using the ChIPBase and TarBase databases, respectively. The ROC curve showed that the expression levels of the six hub genes (*MMP9*, *APOE*, *TNF*, *PPARG*, *CYBA* and *NCF2*) in atherosclerotic samples were accurate in different groups (0.7 < AUC < 0.9). Furthermore, through the analysis of the immune infiltration sclerosis of arterial congestion appearance, the hub genes and immune cells may have a strong positive correlation, prompting the hub genes to release proinflammatory cytokines, further influencing the process of vascular aging and hardening of the arteries. More immune cells may mean more pro-inflammatory cytokines. This observation was further refined by single-cell RNA sequencing (scRNA-seq) analysis of coronary artery disease samples (GSE184073), which revealed that the hub genes are predominantly expressed in monocytes and T cells. Specifically, T cells constituted the largest cell subpopulation, followed by monocytes, suggesting their dominant role in the pathogenic landscape of AS. A phenomenon in which higher levels of inflammatory cytokines in tissues lead to a low-grade, sterile and chronic inflammatory state, defined as immunosenescence or inflamm-aging[57]. In addition to aging, other factors that contribute to inflammation include genetic predisposition, obesity, oxidative stress, changes in intestinal barrier permeability, chronic infections, and immune cell defects[58]. Conversely, interventions that reduce inflammatory markers, such as dietary restriction and exercise, can increase longevity[59, 60]. Therefore, promoting the resolution of inflammation is a potential treatment for alleviating cell aging and organ aging. However, it is still unclear which specific immune cell type plays a decisive role in vascular aging and atherosclerosis.

Identifying effective therapeutic strategies targeting these hub genes is of paramount importance. Our molecular docking analysis explored the interaction between curcumin, a natural polyphenol with known anti-inflammatory and anti-atherosclerotic properties, and the six high-performing hub genes. The results demonstrated strong binding affinities (binding energy < -5 kcal/mol) between curcumin and all target genes, with the most pronounced interaction observed with *PPARG*. *PPARG* is a nuclear receptor that regulates lipid metabolism and inflammation; its activation is generally considered protective in AS. The strong binding of curcumin to *PPARG* suggests that curcumin may exert its therapeutic effects partly by modulating *PPARG* activity, thereby improving lipid profiles and suppressing inflammation. Furthermore, the predicted interactions with *TNF* and *NCF2* imply that curcumin could also directly inhibit pro-inflammatory signaling and oxidative stress. These computational findings provide a molecular basis for future experimental studies to validate curcumin as a targeted therapy for vascular aging-related atherosclerosis.

Despite these promising findings, several limitations must be acknowledged. First, while we validated the hub genes using qRT-PCR, the sample size for experimental validation was relatively small. Larger clinical cohorts are needed to confirm the diagnostic utility of these genes. Second, the bioinformatics analysis relied on publicly available bulk RNA-seq and microarray data, which may mask cell-type-specific nuances that only scRNA-seq can fully resolve; although we incorporated scRNA-seq data for validation, further dedicated single-cell studies on vascular tissues would be beneficial. Third, the molecular docking results are theoretical predictions; *in vitro* and *in vivo* experiments are required to verify the actual binding and functional effects of curcumin on these targets. Finally, the precise mechanistic link between vascular senescence markers and the specific immune subsets (e.g., specific T cell subtypes like Th1 or Th17) warrants deeper investigation using spatial transcriptomics to map these interactions within the plaque architecture.

In conclusion, this study systematically identified a set of vascular aging-related hub genes that are critically involved in the pathogenesis of atherosclerosis. We highlighted the pivotal roles of monocytes and T cells, particularly through *TNF*-mediated inflammation, as key drivers of the disease. The high diagnostic accuracy of these genes offers potential for clinical application, while the predicted interaction with curcumin opens new avenues for therapeutic intervention. By bridging the gap between vascular aging, immune dysregulation, and potential natural therapeutics, our work provides a comprehensive framework for understanding and treating atherosclerosis. Future research should focus on translating these bioinformatics discoveries into clinical biomarkers and targeted therapies to mitigate the burden of this pervasive cardiovascular disease.

## Acknowledgments

The authors would like to thank all staff who made it possible to perform this study.

## Author Contributions

X.F.C., K.L.Z. and J.Y.W. conceived and designed research; X.F.C., K.L.Z. and W.C.W collected data and conducted research; X.F.C., W.C.W.and J.Y.W. analyzed and interpreted data; X.F.C., K.L.Z., and J.Y.W. wrote the initial paper; X.F.C and J.Y.W. revised the paper; All authors read and approved the final manuscript.

## Data Availability Statement

The original contributions presented in the study are included in the article/Supplementary Material, further inquiries can be directed to the corresponding author.

## Supporting information

**S1 Fig. Flowchart of the comprehensive analysis of VARDEGs.** AS, atherosclerosis; GSEA, gene set enrichment analysis; DEGs, differentially expressed genes; VARGs, vascular aging-related genes; GO, gene ontology; KEGG, Kyoto Encyclopedia of Genes and Genomes; VARDEGs, vascular aging-related differentially expressed genes; PPI, protein–protein interaction; ROC, receiver operating characteristic; TF, transcription factor; ssGSEA, single-sample gene-set enrichment analysis.

**S2 Fig. Batch effect removal of GSE100927 and GSE43292.** A. Box plot of combined GEO dataset distribution before batch removal. B. Post-batch integrated GEO datasets (combined datasets) distribution boxplots. C. PCA plot of integrated GEO datasets (combined datasets) before debatching. D. PCA plot of integrated GEO datasets (combined datasets) after debatching. The atherosclerosis datasets GSE100927 and GSE43292 are presented in green and yellow, respectively. GEO, Gene Expression Omnibus; PCA, principal component analysis; AS, atherosclerosis.

**S1 Table. The list of 69 protein-coding vascular aging-related genes (VARGs) obtained from the GeneCards database.**

**S2 Table. The list of 28 vascular aging-related differentially expressed genes (VARDEGs) identified in the combined GEO datasets.**

**S3 Table. The list of 42 transcription factors (TFs) regulating the 6 hub genes identified through the ChIPBase database.**

**S4 Table. The list of 30 microRNAs (miRNAs) associated with the 3 hub genes obtained from the TarBase database.**

**S5 Table. Detailed results of Gene Ontology (GO) and KEGG pathway enrichment analysis for the 28 VARDEGs.**

**S6 Table. Detailed results of Gene Set Enrichment Analysis (GSEA) for the combined GEO datasets.**

## References

1. Ma X, Zhang T, Luo Z, Li X, Lin M, Li R, et al. Functional nano-vector boost anti-atherosclerosis efficacy of berberine in Apoe ((-/-)) mice. Acta Pharm Sin B. 2020;10(9):1769–83. Epub 20200408. doi: 10.1016/j.apsb.2020.03.005. PubMed PMID: 33088695; PubMed Central PMCID: PMCPMC7564017.

2. Bu LL, Yuan HH, Xie LL, Guo MH, Liao DF, Zheng XL. New Dawn for Atherosclerosis: Vascular Endothelial Cell Senescence and Death. Int J Mol Sci. 2023;24(20). Epub 20231013. doi: 10.3390/ijms242015160. PubMed PMID: 37894840; PubMed Central PMCID: PMCPMC10606899.

3. Gimbrone MA, Jr., Garcia-Cardena G. Endothelial Cell Dysfunction and the Pathobiology of Atherosclerosis. Circul Res. 2016;118(4):620–36. doi: 10.1161/CIRCRESAHA.115.306301. PubMed PMID: 26892962; PubMed Central PMCID: PMCPMC4762052.

4. Zhang C, Chen X, Wang JK, Li Y, Cui SJ, Wang Z, et al. Phenotypic Switching of Atherosclerotic Smooth Muscle Cells is Regulated by Activated PARP1-Dependent TET1 Expression. #N/A. 2021;28(7):716–29. Epub 20200925. doi: 10.5551/jat.55343. PubMed PMID: 32981917; PubMed Central PMCID: PMCPMC8265424.

5. Roth Flach RJ, Skoura A, Matevossian A, Danai LV, Zheng W, Cortes C, et al. Endothelial protein kinase MAP4K4 promotes vascular inflammation and atherosclerosis. Nat Commun. 2015;6:8995. Epub 20151221. doi: 10.1038/ncomms9995. PubMed PMID: 26688060; PubMed Central PMCID: PMCPMC4703891.

6. Kong P, Cui ZY, Huang XF, Zhang DD, Guo RJ, Han M. Inflammation and atherosclerosis: signaling pathways and therapeutic intervention. Signal Transduct Target Ther. 2022;7(1):131. Epub 20220422. doi: 10.1038/s41392-022-00955-7. PubMed PMID: 35459215; PubMed Central PMCID: PMCPMC9033871.

7. Dai H, Younis A, Kong JD, Bragazzi NL, Wu J. Trends and Regional Variation in Prevalence of Cardiovascular Risk Factors and Association With Socioeconomic Status in Canada, 2005-2016. JAMA Netw Open. 2021;4(8):e2121443. Epub 20210802. doi: 10.1001/jamanetworkopen.2021.21443. PubMed PMID: 34410395; PubMed Central PMCID: PMCPMC8377569.

8. Xu Z, Han Y, Liu J, Jiang F, Hu H, Wang Y, et al. MiR-135b-5p and MiR-499a-3p Promote Cell Proliferation and Migration in Atherosclerosis by Directly Targeting MEF2C. Sci Rep. 2015;5:12276. Epub 20150717. doi: 10.1038/srep12276. PubMed PMID: 26184978; PubMed Central PMCID: PMCPMC4505325.

9. Wolf D, Ley K. Immunity and Inflammation in Atherosclerosis. Circul Res. 2019;124(2):315–27. doi: 10.1161/CIRCRESAHA.118.313591. PubMed PMID: 30653442; PubMed Central PMCID: PMCPMC6342482.

10. Ungvari Z, Tarantini S, Donato AJ, Galvan V, Csiszar A. Mechanisms of Vascular Aging. Circul Res. 2018;123(7):849–67. doi: 10.1161/CIRCRESAHA.118.311378. PubMed PMID: 30355080; PubMed Central PMCID: PMCPMC6248882.

11. Ungvari Z, Tarantini S, Sorond F, Merkely B, Csiszar A. Mechanisms of Vascular Aging, A Geroscience Perspective: JACC Focus Seminar. J Am Coll Cardiol. 2020;75(8):931–41. doi: 10.1016/j.jacc.2019.11.061. PubMed PMID: 32130929; PubMed Central PMCID: PMCPMC8559983.

12. Sun Y, Wang X, Liu T, Zhu X, Pan X. The multifaceted role of the SASP in atherosclerosis: from mechanisms to therapeutic opportunities. Cell Biosci. 2022;12(1):74. Epub 20220531. doi: 10.1186/s13578-022-00815-5. PubMed PMID: 35642067; PubMed Central PMCID: PMCPMC9153125.

13. Gorgoulis V, Adams PD, Alimonti A, Bennett DC, Bischof O, Bishop C, et al. Cellular Senescence: Defining a Path Forward. Cell. 2019;179(4):813–27. Epub 2019/11/02. doi: 10.1016/j.cell.2019.10.005. PubMed PMID: 31675495.

14. Davis S, Meltzer PS. GEOquery: a bridge between the Gene Expression Omnibus (GEO) and BioConductor. Bioinformatics. 2007;23(14):1846–7. Epub 20070512. doi: 10.1093/bioinformatics/btm254. PubMed PMID: 17496320.

15. Barrett T, Wilhite SE, Ledoux P, Evangelista C, Kim IF, Tomashevsky M, et al. NCBI GEO: archive for functional genomics data sets--update. Nucleic Acids Res. 2013;41(Database issue):D991–5. Epub 20121127. doi: 10.1093/nar/gks1193. PubMed PMID: 23193258; PubMed Central PMCID: PMCPMC3531084.

16. Steenman M, Espitia O, Maurel B, Guyomarch B, Heymann MF, Pistorius MA, et al. Identification of genomic differences among peripheral arterial beds in atherosclerotic and healthy arteries. Sci Rep. 2018;8(1):3940. Epub 20180302. doi: 10.1038/s41598-018-22292-y. PubMed PMID: 29500419; PubMed Central PMCID: PMCPMC5834518.

17. Ayari H, Bricca G. Identification of two genes potentially associated in iron-heme homeostasis in human carotid plaque using microarray analysis. J Biosci (Bangalore). 2013;38(2):311–5. doi: 10.1007/s12038-013-9310-2. PubMed PMID: 23660665.

18. Stelzer G, Rosen N, Plaschkes I, Zimmerman S, Twik M, Fishilevich S, et al. The GeneCards Suite: From Gene Data Mining to Disease Genome Sequence Analyses. Curr Protoc Bioinformatics. 2016;54:1 30 1-1 3. Epub 20160620. doi: 10.1002/cpbi.5. PubMed PMID: 27322403.

19. Leek JT, Johnson WE, Parker HS, Jaffe AE, Storey JD. The sva package for removing batch effects and other unwanted variation in high-throughput experiments. Bioinformatics. 2012;28(6):882–3. Epub 20120117. doi: 10.1093/bioinformatics/bts034. PubMed PMID: 22257669; PubMed Central PMCID: PMCPMC3307112.

20. Ritchie ME, Phipson B, Wu D, Hu Y, Law CW, Shi W, et al. limma powers differential expression analyses for RNA-sequencing and microarray studies. Nucleic Acids Res. 2015;43(7):e47. Epub 20150120. doi: 10.1093/nar/gkv007. PubMed PMID: 25605792; PubMed Central PMCID: PMCPMC4402510.

21. Ben Salem K, Ben Abdelaziz A. Principal Component Analysis (PCA). Tunis Med. 2021;99(4):383–9. PubMed PMID: 35244921; PubMed Central PMCID: PMCPMC8734479.

22. Zhang H, Meltzer P, Davis S. RCircos: an R package for Circos 2D track plots. BMC Bioinformatics. 2013;14:244. Epub 20130810. doi: 10.1186/1471-2105-14-244. PubMed PMID: 23937229; PubMed Central PMCID: PMCPMC3765848.

23. Mi H, Muruganujan A, Ebert D, Huang X, Thomas PD. PANTHER version 14: more genomes, a new PANTHER GO-slim and improvements in enrichment analysis tools. Nucleic Acids Res. 2019;47(D1):D419–D26. doi: 10.1093/nar/gky1038. PubMed PMID: 30407594; PubMed Central PMCID: PMCPMC6323939.

24. Kanehisa M, Goto S. KEGG: kyoto encyclopedia of genes and genomes. Nucleic Acids Res. 2000;28(1):27–30. doi: 10.1093/nar/28.1.27. PubMed PMID: 10592173; PubMed Central PMCID: PMCPMC102409.

25. Yu G, Wang LG, Han Y, He QY. clusterProfiler: an R package for comparing biological themes among gene clusters. #N/A. 2012;16(5):284–7. Epub 20120328. doi: 10.1089/omi.2011.0118. PubMed PMID: 22455463; PubMed Central PMCID: PMCPMC3339379.

26. Subramanian A, Tamayo P, Mootha VK, Mukherjee S, Ebert BL, Gillette MA, et al. Gene set enrichment analysis: a knowledge-based approach for interpreting genome-wide expression profiles. Proc Natl Acad Sci U S A. 2005;102(43):15545–50. Epub 20050930. doi: 10.1073/pnas.0506580102. PubMed PMID: 16199517; PubMed Central PMCID: PMCPMC1239896.

27. Liberzon A, Subramanian A, Pinchback R, Thorvaldsdottir H, Tamayo P, Mesirov JP. Molecular signatures database (MSigDB) 3.0. Bioinformatics. 2011;27(12):1739–40. Epub 20110505. doi: 10.1093/bioinformatics/btr260. PubMed PMID: 21546393; PubMed Central PMCID: PMCPMC3106198.

28. Szklarczyk D, Gable AL, Lyon D, Junge A, Wyder S, Huerta-Cepas J, et al. STRING v11: protein-protein association networks with increased coverage, supporting functional discovery in genome-wide experimental datasets. Nucleic Acids Res. 2019;47(D1):D607–D13. doi: 10.1093/nar/gky1131. PubMed PMID: 30476243; PubMed Central PMCID: PMCPMC6323986.

29. Shannon P, Markiel A, Ozier O, Baliga NS, Wang JT, Ramage D, et al. Cytoscape: a software environment for integrated models of biomolecular interaction networks. Genome Res. 2003;13(11):2498–504. doi: 10.1101/gr.1239303. PubMed PMID: 14597658; PubMed Central PMCID: PMCPMC403769.

30. Yang X, Li Y, Lv R, Qian H, Chen X, Yang CF. Study on the Multitarget Mechanism and Key Active Ingredients of Herba Siegesbeckiae and Volatile Oil against Rheumatoid Arthritis Based on Network Pharmacology. Evid Based Complement Alternat Med. 2019;2019:8957245. Epub 20191126. doi: 10.1155/2019/8957245. PubMed PMID: 31885670; PubMed Central PMCID: PMCPMC6899322.

31. Zhou KR, Liu S, Sun WJ, Zheng LL, Zhou H, Yang JH, et al. ChIPBase v2.0: decoding transcriptional regulatory networks of non-coding RNAs and protein-coding genes from ChIP-seq data. Nucleic Acids Res. 2017;45(D1):D43–D50. Epub 20161023. doi: 10.1093/nar/gkw965. PubMed PMID: 27924033; PubMed Central PMCID: PMCPMC5210649.

32. Li JH, Liu S, Zhou H, Qu LH, Yang JH. starBase v2.0: decoding miRNA-ceRNA, miRNA-ncRNA and protein-RNA interaction networks from large-scale CLIP-Seq data. Nucleic Acids Res. 2014;42(Database issue):D92–7. Epub 20131201. doi: 10.1093/nar/gkt1248. PubMed PMID: 24297251; PubMed Central PMCID: PMCPMC3964941.

33. Xiao B, Liu L, Li A, Xiang C, Wang P, Li H, et al. Identification and Verification of Immune-Related Gene Prognostic Signature Based on ssGSEA for Osteosarcoma. Front Oncol. 2020;10:607622. Epub 20201215. doi: 10.3389/fonc.2020.607622. PubMed PMID: 33384961; PubMed Central PMCID: PMCPMC7771722.

34. Yadav R, Mishra S, Chaturvedi R, Pandey A. Therapeutic potential of curcumin in cardiovascular disease: Targeting atherosclerosis pathophysiology. Biomed Pharmacother. 2025;190:118412. Epub 20250804. doi: 10.1016/j.biopha.2025.118412. PubMed PMID: 40763486.

35. Zhang W, Lv Z, Zhang Y, Gopinath SCB, Yuan Y, Huang D, et al. Targeted Diagnosis, Therapeutic Monitoring, and Assessment of Atherosclerosis Based on Mesoporous Silica Nanoparticles Coated with cRGD-Platelets. Oxid Med Cell Longev. 2022;2022:6006601. Epub 20220929. doi: 10.1155/2022/6006601. PubMed PMID: 36211824; PubMed Central PMCID: PMCPMC9537012.

36. Zhang F, Xia X, Chai R, Xu R, Xu Q, Liu M, et al. Inhibition of USP14 suppresses the formation of foam cell by promoting CD36 degradation. J Cell Mol Med. 2020;24(6):3292–302. Epub 20200122. doi: 10.1111/jcmm.15002. PubMed PMID: 31970862; PubMed Central PMCID: PMCPMC7131911.

37. Jiang ZM, Wu T, Xue YT, Li Y, Li GH, Huang K, et al. Decoding the Mechanism of Shixiao Powder in Treating Coronary Heart Disease Based on Network Pharmacology and Molecular Docking. Evid Based Complement Alternat Med. 2022;2022:3756668. Epub 20220706. doi: 10.1155/2022/3756668. PubMed PMID: 35845584; PubMed Central PMCID: PMCPMC9279019.

38. Ma S, Xie X, Yuan R, Xin Q, Miao Y, Leng SX, et al. Vascular Aging and Atherosclerosis: A Perspective on Aging. Aging Dis. 2024. Epub 20240306. doi: 10.14336/AD.2024.0201-1. PubMed PMID: 38502584.

39. Liberale L, Badimon L, Montecucco F, Luscher TF, Libby P, Camici GG. Inflammation, Aging, and Cardiovascular Disease: JACC Review Topic of the Week. J Am Coll Cardiol. 2022;79(8):837–47. doi: 10.1016/j.jacc.2021.12.017. PubMed PMID: 35210039; PubMed Central PMCID: PMCPMC8881676.

40. Laina A, Stellos K, Stamatelopoulos K. Vascular ageing: Underlying mechanisms and clinical implications. Exp Gerontol. 2018;109:16–30. Epub 2017/06/19. doi: 10.1016/j.exger.2017.06.007. PubMed PMID: 28624356.

41. Stojanovic SD, Fiedler J, Bauersachs J, Thum T, Sedding DG. Senescence-induced inflammation: an important player and key therapeutic target in atherosclerosis. Eur Heart J. 2020;41(31):2983–96. doi: 10.1093/eurheartj/ehz919. PubMed PMID: 31898722; PubMed Central PMCID: PMCPMC7453834.

42. Liu B, Su L, Loo SJ, Gao Y, Khin E, Kong X, et al. Matrix metallopeptidase 9 contributes to the beginning of plaque and is a potential biomarker for the early identification of atherosclerosis in asymptomatic patients with diabetes. Front Endocrinol (Lausanne). 2024;15:1369369. Epub 20240410. doi: 10.3389/fendo.2024.1369369. PubMed PMID: 38660518; PubMed Central PMCID: PMCPMC11039961.

43. Michaud M, Balardy L, Moulis G, Gaudin C, Peyrot C, Vellas B, et al. Proinflammatory cytokines, aging, and age-related diseases. J Am Med Dir Assoc. 2013;14(12):877–82. Epub 20130620. doi: 10.1016/j.jamda.2013.05.009. PubMed PMID: 23792036.

44. Guo S, Mao X, Liu J. Multi-faceted roles of C1q/TNF-related proteins family in atherosclerosis. Front Immunol. 2023;14:1253433. Epub 20231013. doi: 10.3389/fimmu.2023.1253433. PubMed PMID: 37901246; PubMed Central PMCID: PMCPMC10611500.

45. Wang H, Tian Q, Zhang R, Du Q, Hu J, Gao T, et al. Nobiletin alleviates atherosclerosis by inhibiting lipid uptake via the PPARG/CD36 pathway. Lipids Health Dis. 2024;23(1):76. Epub 20240311. doi: 10.1186/s12944-024-02049-5. PubMed PMID: 38468335; PubMed Central PMCID: PMCPMC10926578.

46. Zhang Z, Chen Y, Fu X, Chen L, Wang J, Zheng Q, et al. Identification of PPARG as key gene to link coronary atherosclerosis disease and rheumatoid arthritis via microarray data analysis. PLoS One. 2024;19(4):e0300022. Epub 20240404. doi: 10.1371/journal.pone.0300022. PubMed PMID: 38573982; PubMed Central PMCID: PMCPMC10994321.

47. Balcerzyk-Matic A, Nowak T, Mizia-Stec K, Iwanicka J, Iwanicki T, Banka P, et al. Polymorphic Variants of AGT, ABCA1, and CYBA Genes Influence the Survival of Patients with Coronary Artery Disease: A Prospective Cohort Study. Genes (Basel). 2022;13(11). Epub 20221118. doi: 10.3390/genes13112148. PubMed PMID: 36421822; PubMed Central PMCID: PMCPMC9690336.

48. Huo TM, Wang ZW. Comprehensive Analysis to Identify Key Genes Involved in Advanced Atherosclerosis. Dis Markers. 2021;2021:4026604. Epub 20211210. doi: 10.1155/2021/4026604. PubMed PMID: 34925641; PubMed Central PMCID: PMCPMC8683248.

49. Libby P, Buring JE, Badimon L, Hansson GK, Deanfield J, Bittencourt MS, et al. Atherosclerosis. Nat Rev Dis Primers. 2019;5(1):56. Epub 20190816. doi: 10.1038/s41572-019-0106-z. PubMed PMID: 31420554.

50. Pan H, Ho SE, Xue C, Cui J, Johanson QS, Sachs N, et al. Atherosclerosis Is a Smooth Muscle Cell-Driven Tumor-Like Disease. Circulation. 2024;149(24):1885–98. Epub 20240430. doi: 10.1161/CIRCULATIONAHA.123.067587. PubMed PMID: 38686559; PubMed Central PMCID: PMCPMC11164647.

51. Henein MY, Vancheri S, Longo G, Vancheri F. The Role of Inflammation in Cardiovascular Disease. Int J Mol Sci. 2022;23(21). Epub 20221026. doi: 10.3390/ijms232112906. PubMed PMID: 36361701; PubMed Central PMCID: PMCPMC9658900.

52. Ajay AK, Zhu LJ, Zhao L, Liu Q, Ding Y, Chang YC, et al. Local vascular Klotho mediates diabetes-induced atherosclerosis via ERK1/2 and PI3-kinase-dependent signaling pathways. Atherosclerosis. 2024;396:118531. Epub 20240703. doi: 10.1016/j.atherosclerosis.2024.118531. PubMed PMID: 38996716.

53. Kong X, Chen H, Li D, Ma L. Effects of imbalance of lipid metabolism through NF-KB pathway on atherosclerosis and vascular aging in rats. Cell Mol Biol (Noisy-le-grand). 2022;67(5):144–50. Epub 20220204. doi: 10.14715/cmb/2021.67.5.20. PubMed PMID: 35818259.

54. Fu X, Sun Z, Long Q, Tan W, Ding H, Liu X, et al. Glycosides from Buyang Huanwu Decoction inhibit atherosclerotic inflammation via JAK/STAT signaling pathway. Phytomedicine. 2022;105:154385. Epub 20220808. doi: 10.1016/j.phymed.2022.154385. PubMed PMID: 35987015.

55. Hao L, Ren M, Rong B, Xie F, Lin MJ, Zhao YC, et al. TWEAK/Fn14 mediates atrial-derived HL-1 myocytes hypertrophy via JAK2/STAT3 signalling pathway. J Cell Mol Med. 2018;22(9):4344–53. Epub 20180704. doi: 10.1111/jcmm.13724. PubMed PMID: 29971943; PubMed Central PMCID: PMCPMC6111870.

56. Chen M, Xiao L, Dai G, Lu P, Zhang Y, Li Y, et al. Inhibition of JAK-STAT Signaling Pathway Alleviates Age-Related Phenotypes in Tendon Stem/Progenitor Cells. Front Cell Dev Biol. 2021;9:650250. Epub 20210329. doi: 10.3389/fcell.2021.650250. PubMed PMID: 33855026; PubMed Central PMCID: PMCPMC8039155.

57. Franceschi C, Bonafe M, Valensin S, Olivieri F, De Luca M, Ottaviani E, et al. Inflamm-aging. An evolutionary perspective on immunosenescence. Ann N Y Acad Sci. 2000;908:244–54. doi: 10.1111/j.1749-6632.2000.tb06651.x. PubMed PMID: 10911963.

58. Rodier F, Coppe JP, Patil CK, Hoeijmakers WA, Munoz DP, Raza SR, et al. Persistent DNA damage signalling triggers senescence-associated inflammatory cytokine secretion. Nat Cell Biol. 2009;11(8):973–9. Epub 20090713. doi: 10.1038/ncb1909. PubMed PMID: 19597488; PubMed Central PMCID: PMCPMC2743561.

59. Barzilai N, Huffman DM, Muzumdar RH, Bartke A. The critical role of metabolic pathways in aging. Diabetes. 2012;61(6):1315–22. doi: 10.2337/db11-1300. PubMed PMID: 22618766; PubMed Central PMCID: PMCPMC3357299.

60. Fontana L. Neuroendocrine factors in the regulation of inflammation: excessive adiposity and calorie restriction. Exp Gerontol. 2009;44(1-2):41–5. Epub 20080412. doi: 10.1016/j.exger.2008.04.005. PubMed PMID: 18502597; PubMed Central PMCID: PMCPMC2652518.

